# Prior selfing occur in larger flowers in a mixed mating species with delayed selfing

**DOI:** 10.1101/2021.12.08.471669

**Authors:** Kana Masuda, Atushi Ushimaru

## Abstract

Theory predicts that prior self-pollination (prior selfing) should not evolve in mixed mating species that enable delayed selfing. In this study, we test the hypotheais that prior selfing has evolved under severe pollinator limitation in the mixed mating species *Commelina communis* which can reproduce via delayed selfing. The hypothesis predicts that prior selfing occurs more frequently in populations with very low pollinator availability and/or in smaller flowers which receive infrequent visitations. We tested the predictions by comparing the degree of prior selfing among ten populations experiencing various levels of pollinator limitation and by examining a relationship between individual flower size and the occurrence of prior selfing. Populations with higher pollinator availability had higher prior selfing rate. Moreover, prior selfing occurs more frequently in larger flowers. These findings were totally opposite patterns of the predictions and the previous findings. We proposed new hypotheses that prior selfing has been maintained by the presence of reproductive interference from the congener and/or propotency in *C. communis* to explain our unexpected findings. We should verify potential effects of reproductive interference and propotency in future to elucidate the mystery of prior selfing in this mixed mating species with delayed selfing.

---

In mixed mating species, which exhibit both outcrossing and selfing, prior self-pollination (prior selfing) is less likely to evolve than delayed one (Lloyd 1992). This is because delayed selfing that occurs at flower ending stage does not diminish opportunities for outcrossing and provides reproductive assurance when pollinator visits are limited (Becerra and Lloyd 1992, Lloyd 1992, Kalisz and Vogler 2003) whereas prior selfing can cause seed discounting which diminish chances to outcross (Lloyd 1992). Therefore, prior selfing should not be found in mixed mating species that enable delayed selfing (Lloyd 1992).

Form this evolutionary point of view, *Commelina communis* is a mysterious species, because this mixed mating species reproduces partly via prior selfing by means of bud pollination as well as via delayed selfing (Morita and Nigorikawa 1999, Katsuhara and Ushimaru 2019). The evolutionary conditions for prior selfing are stringent, such that it would be selected only under strong pollinator limitation or conditions preferring complete selfing owing to no or very low inbreeding depression within a population (Lloyd 1992, Spigler and Kalisz 2017). *Commelina communis* is presumably facultatively xenogamous by the presence of staminate flowers (andromonoecy), the high pollen:ovule ratio and pollinator-mediated pollen transfer (Ushimaru et al. 2014), thus the selection favoring complete selfing unlikely exists. Thus, the evolution of prior selfing might have occurred in pollinator-limited populations. Indeed, *C. communis* populations often suffered from severe pollinator limitation (Ushimaru et al. 2014). Otherwise, prior selfing may occur in plants with smaller flowers because small flowers are usually much less frequently visited by pollinators within a population and thus have no or little chances to outcross (Elle and Carney 2003).

In this study, we tested these hypotheses by comparing the degree of prior selfing among ten populations experiencing various levels of pollinator limitation and by examining a relationship between individual flower size and the existence of prior selfing. We predict that prior selfing occurs more frequently in populations with very low pollinator availability and/or in smaller flowers. Based on our results, we discuss the adaptive significance of prior selfing in the mixed mating species enabling delayed selfing.

*Commelina communis* is an annual, andromonoecious herb, natively distributed in temperate north-eastern Asia and often growing around paddy fields and along roadsides. A plant usually produces multiple inflorescences with perfect and/or staminate flowers. Mostly, one flower per inflorescence opens at sunrise and remain open until noon. *Commelina* flowers are heterantherous and have two long pollinating stamens one medium-length rewarding stamen, and three short staminoides. Fertile pollen is produced only on anthers of the long (L-anther) and medium-length (M-anther) stamens produce (Ushimaru et al. 2007). Syrphid flies and bees are main visitors to the flowers of *C. communis* in many fields (Ushimaru et al. 2007, 2009, 2014, Katsuhara and Ushimaru 2019) and their visits are significantly lower in flowers with smaller petals (Murakami et al. unpublished data, see uploaded submitted manuscript).

We transplanted 130 *C. communis* seedlings of from ten populations in a greenhouse without pollinators at Kobe University: 10 seedlings from each of three (pops. 3, 5 and 10) and four populations (pops. 1, 4, 6 and 9) in 2020 and 2021, respectively and those of three populations (pops. 2, 7 and 8) were examined in both years (Appendix S1: Table S1, Fig. S1). We preliminarily examined pollinator availability of the ten populations (Appendix S1: Table S1; Ushimaru et al. unpublished data). We planted and cultivated these seedlings with water every day and fertilizer once every two weeks in the greenhouse.

During the flowering peak from August to September, 2020 and 2021, we arbitrarily selected 172 and 623 (totally 802) newly opened perfect flowers, respectively (Appendix S1: Table S1). Floral trait measurements were conducted together with checking the occurrence of prior selfing. The occurrence of prior selfing was largely depending on the presence of the anther dehiscence before flower opening (Fig. 1, Appendix S2). We measured the heights of the stigma and L-anther using digital calipers and then removed one large blue petal from each flower, flattened it under microscope slides, and measured its length. When possible, we measured the right-hand L-anther and petal from each flower. Degree of anther-stigma separation (herkogamy) was calculated as the stigma height minus the L-anther height. After the measurements, we removed all types of stamens to prevent delayed selfing, thus seeds can be produced only via prior selfing. We counted seeds produced for each flower to calculate prior selfing rate as seed set (the number of mature seeds divided by ovule number, four) three weeks after the treatment. Hand-pollination experiment was conducted to analyze the effects of flower size on resource limitation at seed production in fully pollinated flowers. During the flowering peak in August to September, 2021, we arbitrarily selected 152 newly opened perfect flowers from seven populations (Appendix S1: Table S1). We measured the four floral traits and applied self pollen to the stigma after opening for each flower. We examined seed set for these flowers three weeks after the treatment.

**Fig 1.**
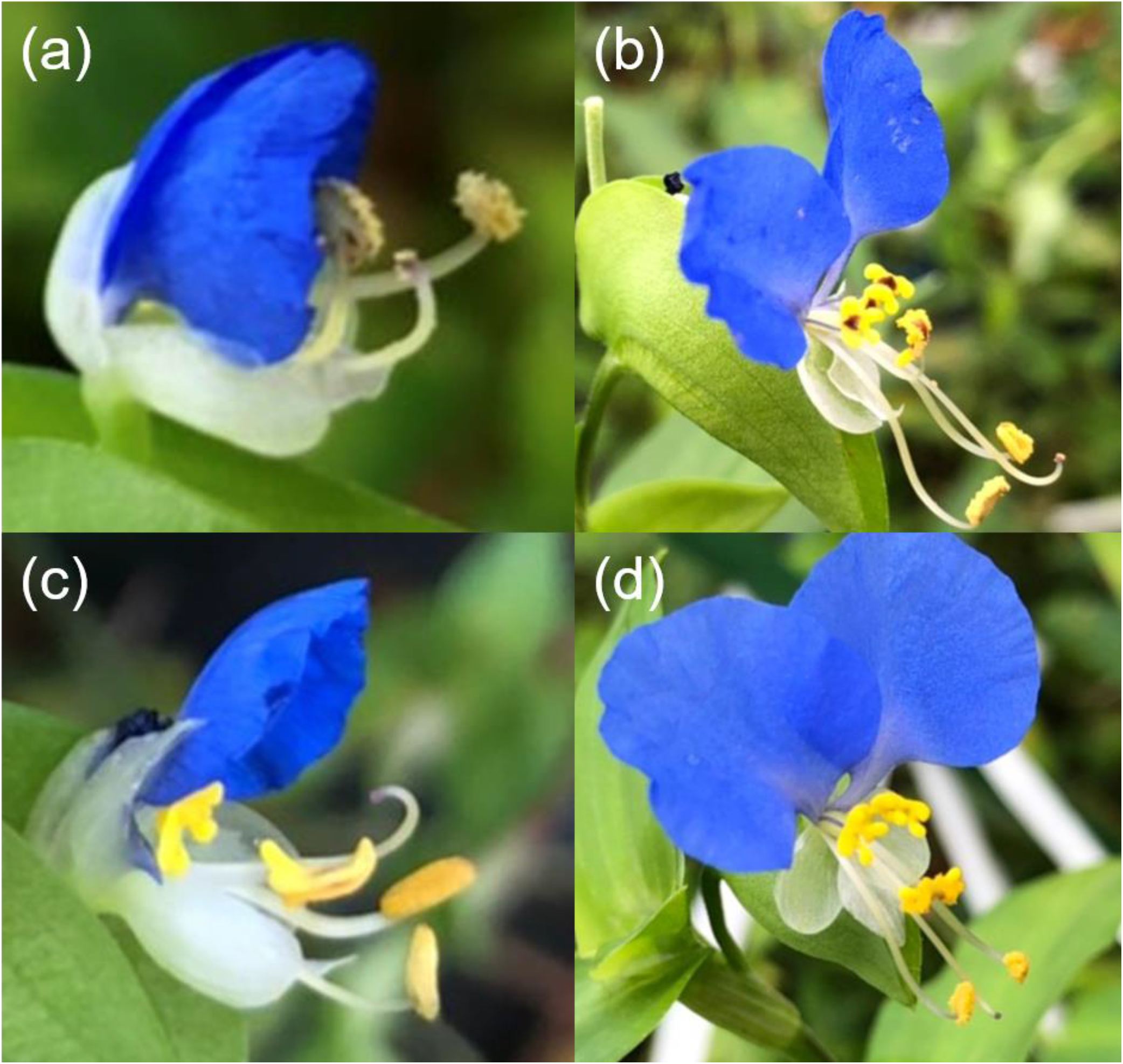
Photographs of the occurrences of anther dehiscence just before opening (a), autonomous prior self-pollination after opening in an anther dehisced flower (b), a flower without the dehisced anthers before opening (c) and unpollinated flower after opening (d).

We firstly examined inter-population differences in the prior selfing rate and the relationship between the population mean pollinator visit frequency (per plot and per flower) and the prior selfing rate using generalized linear mixed models (GLMMs) with binomial errors and logit link and a post hoc test (Tukey-Kramer method). In the GLMM, the prior selfing rate was the response variable whereas population identity or pollinator visit frequency (per plot or per flower) was the explanatory variable and individual identity or nested population and individual identities were random terms, respectively. The significance of each explanatory variable in these and following GLMMs was examined using a Wald test. We performed all the analyses using the glmmADMB package of R (ver. 3.6.1; R Development Core Team; Fournier et al. 2012). We found significant inter-population differences in the prior selfing rate, but the variable did not vary with the pollinator availability (Appendix S1: Table S3, Fig. S2, S4).

We next examined the relationship between each floral measurement (petal length, stigma and L-anther heights, or the degree of herkogamy) and the occurrence of prior selfing separately. A principal component analysis was then performed on the entire floral trait data (i.e., the petal length and stigma and L-anther heights of the ten populations) in order to summarize floral characteristics. The first PC axis (PC1) accounted for 73% of the total variance and positively correlated with all the measurements, thus could be used as the size of each flower (Appendix S1: Table S2). We constructed GLMMs (binomial errors and logit link) in which each floral trait or PC1 and the prior selfing rate were the explanatory and response variables, respectively and the observation date and nested population and individual identities were the random terms. We also analyzed data of each population separately using GLMMs with the observation date and individual identity were the random terms.

We found significant inter-population differences in PC1 but the value was not related with the pollinator visit frequency (Appendix S1: Table S3, Fig. S3, S4). The prior selfing rate significantly increased with increasing the sizes of three floral traits and PC1 (Fig. 2a, Appendix S1: Table S3, Fig. S5). Meanwhile the variable did not vary with the degree of herkogamy (Fig. 2b, Appendix S1: Table S3). Flower size did not affect seed set in artificially-pollinated flowers, indicating no resource limitation in smaller flowers (Appendix S1: Table S3, Fig. S6). Significantly positive relationships between the prior selfing rate and PC1 were found in five populations (Appendix S1: Table S3, Fig. S7). The prior selfing rate significantly increased with the PC1 score in five populations (pops. 1, 2, 3, 5, 8), most of which were those with limited pollinator visits, whereas the rate did not vary with PC1 score in the remaining five populations (Appendix S1: Table S3, Fig. S7).

**Fig 2.**
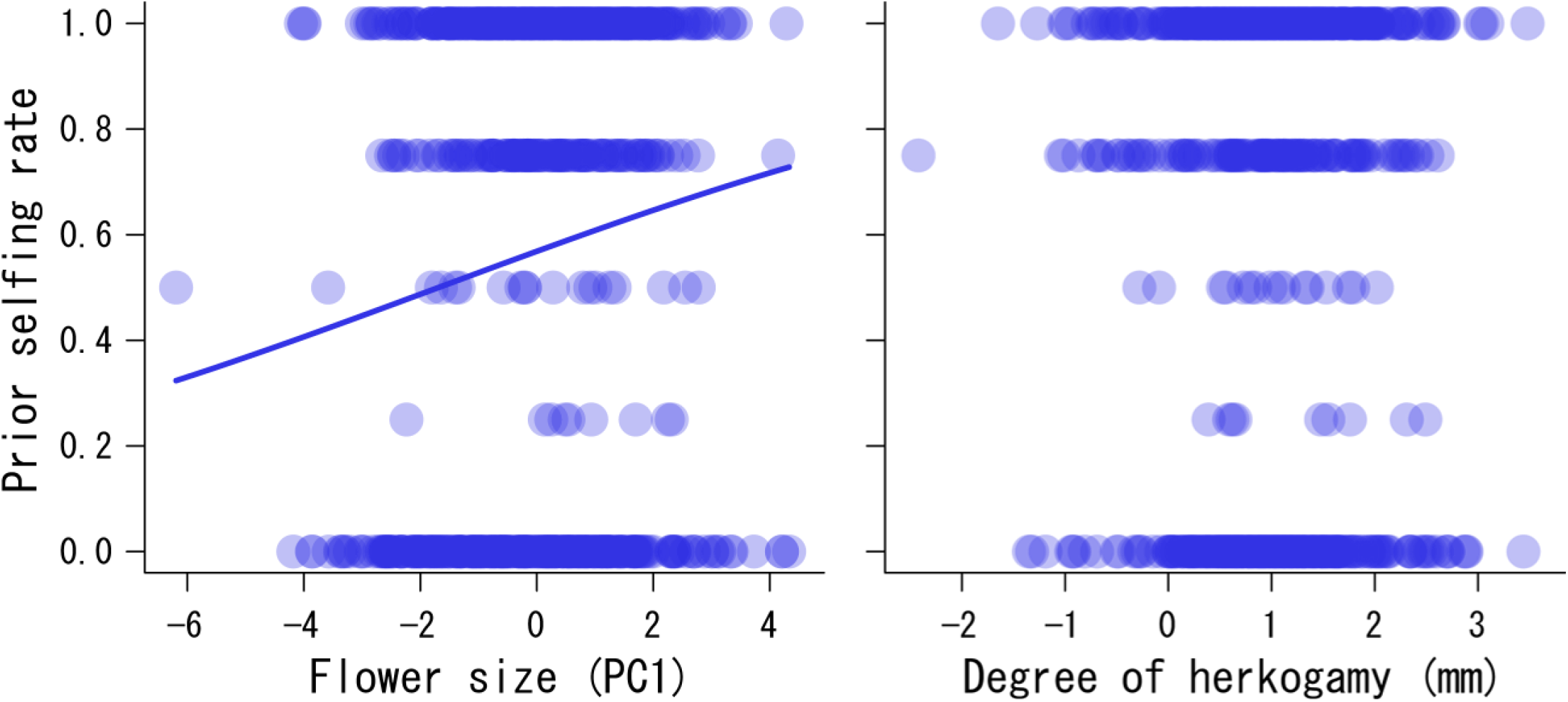
Relationships between flower size (PC1, a) and the degree of herkogamy (b) and the prior selfing rate. Significant relationship is indicated with regression line drawn using estimated coefficients from the GLMM.

In *C. communis*, the population with the highest pollinator availability (pop10) has the highest prior selfing rate, and there was no negative correlation between prior selfing rate and pollinator availability (Appendix S1: Table S3, Fig. S2, S4), giving no support for our prediction and the theory (Lloyd 1992). Prior selfing occurs more frequently in larger flowers, which was totally opposite pattern of the prediction and the previous study (Fig. 2a, Appendix S1: Table S3, Fig. S5; Elle and Carney 2003). The findings contradict the hypothesis that prior selfing evolves to assure reproduction under severe pollinator limitation. Furthermore, we found no relationship between the degree of herkogamy and prior selfing rate (Fig. 2b, Appendix S1: Table S3), being inconsistent with the previous studies (Takebayashi et al. 2006). In *C. communis*, the distance between the anthers and stigma after opening unlikely influence the occurrence of autonomous self-pollination because prior selfing occurs by the dehisced anther contact with the stigma in a bud.

The following hypotheses may explain these unexpected findings. In rural area, *C. communis* often coexists with and suffers from reproductive interference (seed set reduction by heterospecific pollen deposition) from congeneric *C. communis* f. *ciliate* (Appendix S1: Table S1; Katsuhara and Ushimaru 2019). Prior selfing may evolve under reproductive interference by coexisting relatives to mitigate the negative impact (Katsuhara et al. 2021). Rural population with high pollinator availability has potential opportunities to receive heterospecific pollen that reduce seed production. Higher prior selfing rate irrespective of flower size in the rural populations visited by abundant pollinators (pops. 7, 8 and 9) would be adaptation to interspecific competition with the closely-related species.

Meanwhile, higher prior selfing rate in larger flowers can be explained by the idea that these flowers have longer styles in which outcross pollen may overtake self pollen after flower opening. Prior selfing is as advantageous as delayed selfing when the propotency of outcross pollen is strong (Lloyd 1992). Actually, prior selfing takes several hours to fertilize the ovules in *C. communis* (Appendix S3: Fig. S1), likely giving chance of being overtaken by outcross pollen carried by pollinators just after flower opening. Since the effects of reproductive interference from the congener and propotency on the evolution of prior selfing in *C. communis* are just hypotheses, we should verify the potential effects in future to elucidate the mystery of prior selfing in the mixed mating species with delayed selfing.

## Supporting information

https://drive.google.com/file/d/1ugd2gQJLPj2HPQ4fUboyqwbbkV4f_RDL/view?usp=sharing

## Acknowledgements

We thank Kana Murakami and Koki R. Katsuhara for allowing us to use pollinator data and the members of the Biodiversity Laboratory at Kobe University for assistance in data collection. We also thank the members of the Biodiversity Laboratory and Evolutionary Ecology Laboratory at Kobe University for their valuable comments on our study.

## Literature Cited

Becerra, J. X., and Lloyd, D. G. 1992. Competition-dependent abscission of self-pollinated flowers of Phormium tenax (Agavaceae): a second action of self-incompatibility at the whole flower level? Evolution 46: 458–469.

Cruden, R. W. 1977. Pollen-Ovule ratios: A conservative indicator of breeding systems in flowering plants. Evolution 31: 32–46.

Elle, E., and Carney, R. 2003. Reproductive assurance varies with flower size in Collinsia parviflora (Scrophulariaceae). American Journal of Botany 90: 888–896.

Fournier, D. A. et al. 2012. AD Model Builder: using automatic differentiation for statistical inference of highly parameterized complex nonlinear models. Optimization Methods and Software 27: 233–249.

Kalisz, S., and Vogler, D. W. 2003. Benefits of autonomous selfing under unpredictable pollinator environments. Ecology 84: 2928–2942.

Katsuhara, K. R., Tachiki, Y., Iritani, R., and Ushimaru, A. 2021. The eco-evolutionary dynamics of prior selfing rates promote coexistence without niche partitioning under conditions of reproductive interference. Journal of Ecology 109: 3916–3928.

Katsuhara, K. R., and Ushimaru, A. 2019. Prior selfing can mitigate the negative effects of mutual reproductive interference between coexisting congeners. Functional Ecology 33: 1504–1513.

Lloyd, D. G. 1992. Self-and cross-fertilization in plants. II. The selection of self-fertilization. International journal of plant sciences 153: 370–380.

Morita, T., and Nigorikawa, T. 1999. Phenotypic plasticity of floral sex. Pages 227–242 in M. Ohara, ed. Natural history of flowers. Hokkaido University Press, Sapporo [in Japanese]

Spigler, R. B., and Kalisz, S. 2017. Persistent pollinators and the evolution of complete selfing. American journal of botany 104: 1783–1786.

Takebayashi, N., D. E. Wolf, and L.F. 2006. Delph Effect of variation in herkogamy on outcrossing within a population of Gilia achilleifolia. Heredity 96: 159–165.

Ushimaru, A., Dohzono, I., Takami, Y., and Hyodo, F. 2009. Flower orientation enhances pollen transfer in bilaterally symmetrical flowers. Oecologia 160: 667–674.

Ushimaru, A., Kobayashi, A., and Dohzono, I. 2014. Does urbanization promote floral diversification? Implications from changes in herkogamy with pollinator availability in an urban-rural area. The American Naturalist 184: 258–267.

Ushimaru, A., and Hyodo, F. 2005. Why do bilaterally symmetrical flowers orient vertically? Flower orientation influences pollinator landing behaviour. Evolutionary Ecology Research 7: 151–160.

Ushimaru, A., Watanabe, T., and Nakata, K. 2007. Colored floral organs influence pollinator behavior and pollen transfer in Commelina communis (Commelinaceae). American Journal of Botany 94: 249–258.

