## Supplementary material for "Prior selfing occur in larger flowers in a mixed mating species with delayed selfing": https://drive.google.com/file/d/1ugd2gQJLPj2HPQ4fUboyqwbbkV4f_RDL/view?usp=sharing

**Appendix S1.**

Table S1. Study site (population) and measurement description: location, site type, presence of a coexisting congeneric species (*Commelina communis* f. *ciliate*), pollinator visit frequency (PV) per plot and per flower, the number of sample flowers (individuals) for examining prior selfing and hand self-pollination in 2020 and 2021, and the population mean prior selfing rate (PSR) and PC1 score. The numbers of sampled flowers and seedlings, *N* (seedlings), in 2020 and 2021were shown. Ten seedlings for each population were transplanted and grown in a greenhouse in each year.

| | | | | | | | | Sample size | | | | |
|---|---|---|---|---|---|---|---|---|---|---|---|---|
| Location | | | | | | PV* | | PR | | HP | | |
| Population | ID | Latitude | Longitude | Site type | Congener (yes/no) | (/45min/ plot) | (/45min/ flower) | in 2020 | in 2021 | in 2021 | PSR | PC1 |
| Ikoma | 1 | 34.66783 | 135.68135 | Rural | no | 1.167 | 0.016 | | 81 (10) | 20 | 0.556 | 0.195 |
| Turumiryokuchi | 2 | 34.71722 | 135.57683 | Urban | no | 1.667 | 0.05 | 25 (10) | 72 (10) | 25 | 0.582 | -0.023 |
| Fukae | 3 | 34.71983 | 135.29015 | Urban | no | 2.889 | 0.153 | 37 (10) | | | 0.493 | -1.000 |
| Nakacho | 4 | 34.84722 | 134.94318 | Urban | no | 4.778 | 0.118 | | 82 (10) | 21 | 0.494 | 0.642 |
| Maya | 5 | 34.70495 | 135.22656 | Urban | no | 5.111 | 0.1 | 21 (10) | | | 0.512 | -1.618 |
| Karato | 6 | 34.78656 | 135.21858 | Sub-urban | no | 6.000 | 0.107 | | 81 (10) | 17 | 0.525 | 0.986 |
| Byoubu | 7 | 34.81791 | 135.16772 | Rural | yes | 7.111 | 0.182 | 32 (10) | 98 (10) | 19 | 0.644 | -0.081 |
| Sakae | 8 | 34.85912 | 134.99306 | Rural | yes | 7.667 | 0.14 | 40 (10) | 118 (10) | 32 | 0.600 | -0.294 |

| | | | | | | | | | | | | |
|---|---|---|---|---|---|---|---|---|---|---|---|---|
| Yamatanaka | 9 | 34.77515 | 135.1369 | Rural | yes | 8.167 | 0.074 | | 91 (10) | 19 | 0.390 | 0.034 |
| Ogo | 10 | 34.81825 | 135.19345 | Rural | yes | 24.583 | 0.173 | 24 (10) | | | 0.677 | -0.881 |

*In 2018, 2019, we observed pollinator visitations for 2, 3 or 4 days during the peak of anthesis at each study site. We set three 1x1 $m^2$ plots on first observation day at each study site. On each observation day, we recorded all pollinator individuals that visited flowers for 45-min (three 15-min observations) in each plot during anthesis (06:00-10:00). We divided pollinators into five groups: *E. balteatus*, other small syrphid flies, *B. diversus diversus*, *Apis mellifera* and small-sized bees. After pollinator observations, we counted all flowers, including both the perfect and male flowers in each plot.

Table S2. Scores of the compounds on the Principal Components of the PCA performed on floral traits.

| Parameter | PC1 | PC2 | PC3 |
|---|---|---|---|
| Standard deviation | 1.480 | 0.779 | 0.449 |
| Proportion of Variance | 0.730 | 0.202 | 0.067 |
| Floral traits | | | |
| Petal length | 0.499 | 0.866 | 0.041 |
| Stigma height | 0.617 | -0.321 | -0.719 |
| L-anther height | 0.609 | -0.384 | 0.694 |

Table S3. Results of generalised linear mixed model (GLMM) analyses. Analyses of floral trait measurements, hand-pollination treatment, floral trait measurements for each population and timing, mechanism of prior self-pollination and pollinator availability were shown (see text for details of the explanatory variables). Boldface indicates significant effects ($P < 0.05$).

| Response variable/Explanatory variable | Estimated coefficient | Standard error | *z* value | *P* |
|---|---|---|---|---|
| Floral trait measurements | | | | |
| Prior selfing rate/ | | | | |
| Intercept | 0.228 | 0.257 | 0.89 | 0.290 |
| **PC1** | 0.164 | 0.035 | 4.66 | **<0.001** |
| Number of days from August 1 | 0.001 | 0.003 | 0.37 | 0.690 |
| Prior selfing rate/ | | | | |
| **Intercept** | -1.349 | 0.530 | -2.55 | **0.011** |
| **Petal length** | 0.136 | 0.035 | 3.92 | **<0.001** |
| Number of days from August 1 | -0.001 | 0.003 | -0.18 | 0.859 |
| Prior selfing rate/ | | | | |
| **Intercept** | -1.368 | 0.637 | -2.15 | **0.032** |
| **Stigma height** | 0.129 | 0.043 | 3.03 | **0.002** |
| Number of days from August 1 | 0.000 | 0.003 | -0.07 | 0.942 |
| Prior selfing rate/ | | | | |
| **Intercept** | -2.090 | 0.584 | -3.58 | **<0.001** |
| **L-anther height** | 0.207 | 0.045 | 4.63 | **<0.001** |
| Number of days from August 1 | 0.001 | 0.003 | -0.28 | 0.777 |
| Prior selfing rate/ | | | | |

| | | | | |
|---|---|---|---|---|
| Intercept | 0.588 | 0.319 | 1.85 | 0.065 |
| Degree of herkogamy | -0.096 | 0.060 | -1.61 | 0.108 |
| Number of days from August 1 | -0.007 | 0.003 | -2.01 | 0.044 |
| Hand-pollination experiment | | | | |
| Seed set/ | | | | |
| **Intercept** | 8.881 | 3.337 | 2.66 | **0.008** |
| PC1 | -0.036 | 0.136 | -0.27 | 0.790 |
| Number of days from August 1 | -0.564 | 0.296 | -1.91 | 0.056 |
| Floral trait measurements for each population | | | | |
| 1 Prior selfing rate/ | | | | |
| Intercept | 0.120 | 0.457 | 0.26 | 0.793 |
| **PC1** | 0.250 | 0.095 | 2.64 | **0.008** |
| Number of days from August 1 | 0.001 | 0.011 | 0.13 | 0.894 |
| 2 Prior selfing rate/ | | | | |
| Intercept | 0.619 | 0.481 | 1.00 | 0.200 |
| **PC1** | 0.426 | 0.104 | 4.11 | **<0.001** |
| Number of days from August1 | -0.009 | 0.009 | -1.00 | 0.320 |
| 3 Prior selfing rate/ | | | | |
| Intercept | 2.004 | 1.394 | 1.44 | 0.150 |
| **PC1** | 0.346 | 0.160 | 2.16 | **0.030** |
| Number of days from August 1 | -0.045 | 0.030 | -1.52 | 0.130 |
| 4 Prior selfing rate/ | | | | |

| | | | | |
|---|---|---|---|---|
| **Intercept** | 1.186 | 0.482 | 2.46 | **0.014** |
| PC1 | -0.060 | 0.104 | -0.57 | 0.566 |
| **Number of days from August 1** | -0.030 | 0.010 | -2.94 | **0.003** |
| 5 Prior selfing rate/ | | | | |
| Intercept | -4.365 | 3.748 | -1.16 | 0.244 |
| **PC1** | 0.986 | 0.392 | 2.52 | **0.012** |
| Number of days from August 1 | 0.169 | 0.105 | 1.60 | 0.109 |
| 6 Prior selfing rate/ | | | | |
| Intercept | -0.325 | 0.418 | -0.67 | 0.500 |
| PC1 | 0.055 | 0.110 | 0.50 | 0.620 |
| Number of days from August 1 | 0.011 | 0.010 | 1.07 | 0.280 |
| 7 Prior selfing rate/ | | | | |
| Intercept | 0.330 | 0.373 | 0.89 | 0.375 |
| PC1 | 0.089 | 0.087 | 1.03 | 0.302 |
| Number of days from August 1 | 0.013 | 0.008 | 1.67 | 0.095 |
| 8 Prior selfing rate/ | | | | |
| Intercept | 0.122 | 0.373 | 0.33 | 0.744 |
| **PC1** | 0.176 | 0.076 | 2.30 | **0.021** |
| Number of days from August 1 | 0.008 | 0.008 | 0.95 | 0.341 |
| 9 Prior selfing rate/ | | | | |
| Intercept | 0.359 | 0.590 | 0.61 | 0.540 |
| PC1 | -0.024 | 0.161 | -0.15 | 0.880 |
| Number of days from August 1 | -0.018 | 0.012 | -1.54 | 0.120 |
| 10 Prior selfing rate/ | | | | |

| | | | | |
|---|---|---|---|---|
| **Intercept** | -3.899 | 1.493 | -2.61 | **0.009** |
| PC1 | -0.02 | 0.191 | -0.10 | 0.917 |
| **Number of days from August 1** | 0.108 | 0.035 | 3.10 | **0.002** |
| Timing and mechanism of prior self-pollination | | | | |
| The occurrence/absence of autonomous self-pollination after flower opening (1/0)/ | | | | |
| **Intercept** | -3.068 | 0.723 | -4.24 | **<0.001** |
| **The occurrence/absence of anther dehiscence before flower opening (1/0)** | 4.860 | 0.870 | 5.59 | **<0.001** |
| Pollinator availability | | | | |
| PC1/ | | | | |
| Intercept | -0.195 | 0.501 | -0.39 | 0.700 |
| Pollinator visit frequency (45min/plot) | -0.002 | 0.022 | -0.09 | 0.930 |
| PC1/ | | | | |
| Intercept | -0.256 | 0.557 | -0.46 | 0.650 |
| Pollinator visit frequency (45min/flower) | 0.424 | 2.417 | 0.18 | 0.860 |
| Prior selfing rate/ | | | | |
| Intercept | 0.129 | 0.220 | 0.59 | 0.560 |
| Pollinator visit frequency (45min/plot) | 0.024 | 0.023 | 1.01 | 0.310 |
| Prior selfing rate/ | | | | |
| Intercept | -0.140 | 0.305 | -0.46 | 0.650 |
| Pollinator visit frequency (45min/flower) | 3.725 | 2.327 | 1.60 | 0.110 |

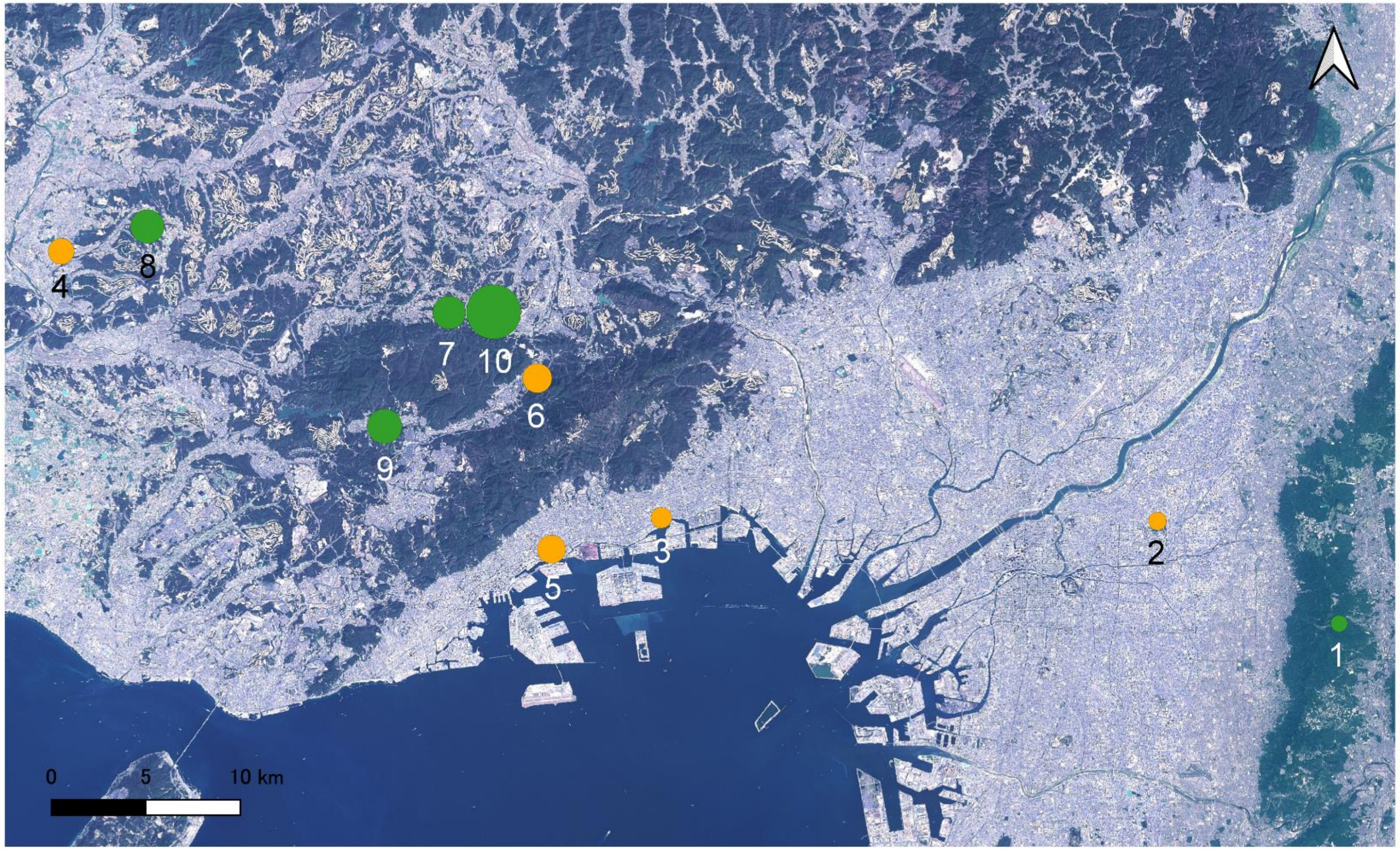

Fig S1. Map of the 10 study sites in the Osaka-Kobe area. Four urban (2-5), one suburban (6), and five rural (1, 7-10) populations were examined. Circle size indicates pollinator visit frequency (/45min/plot).

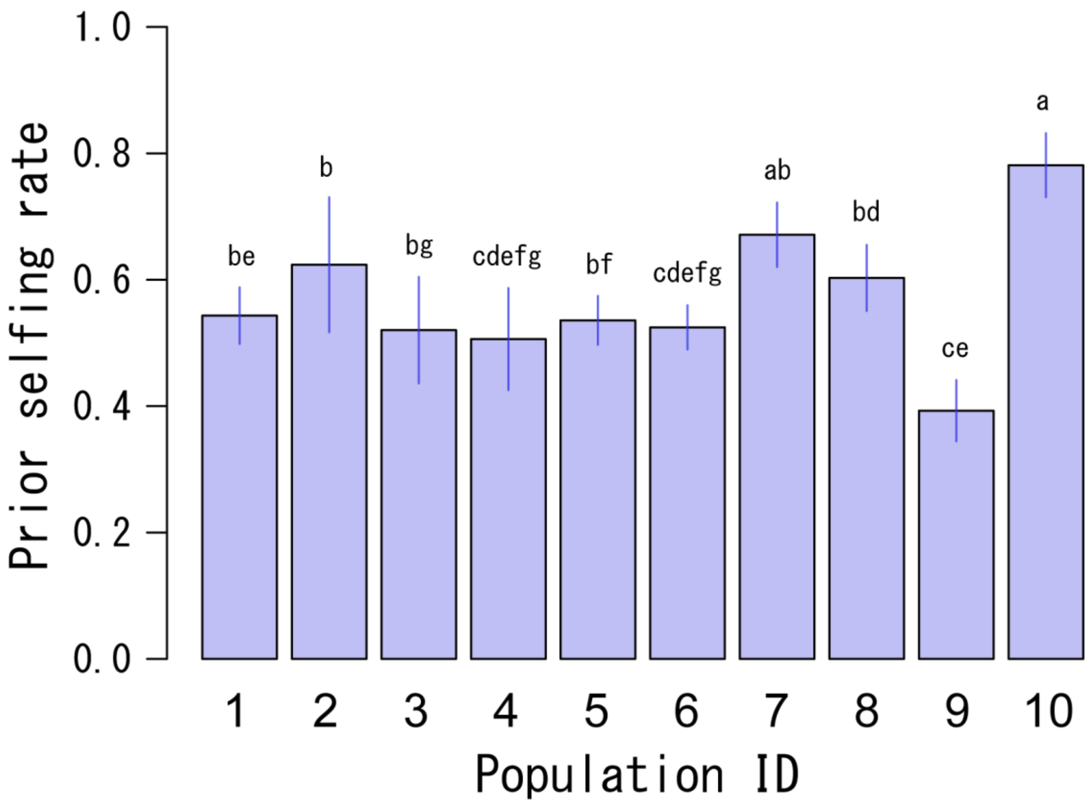

Fig S2. Prior selfing rate (mean ± SE) via prior selfing in ten populations. Different letters above bars indicate significant ($p < 0.05$) differences between populations based on Tukey-Kramer method.

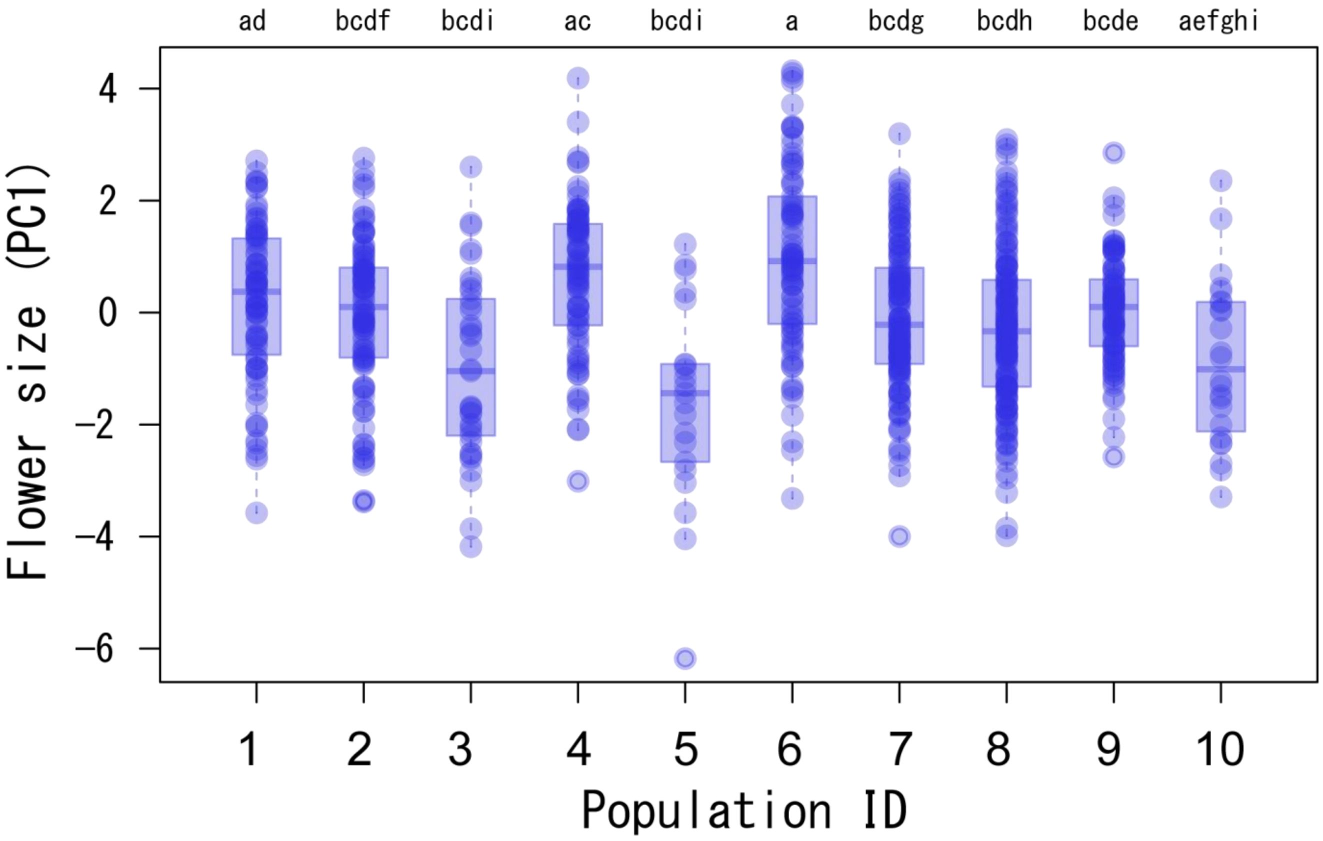

Fig S3. Flower size (PC1) in ten populations. Different letters above bars indicate significant ($p < 0.05$) differences between populations based on the GLMM and Tukey-Kramer method.

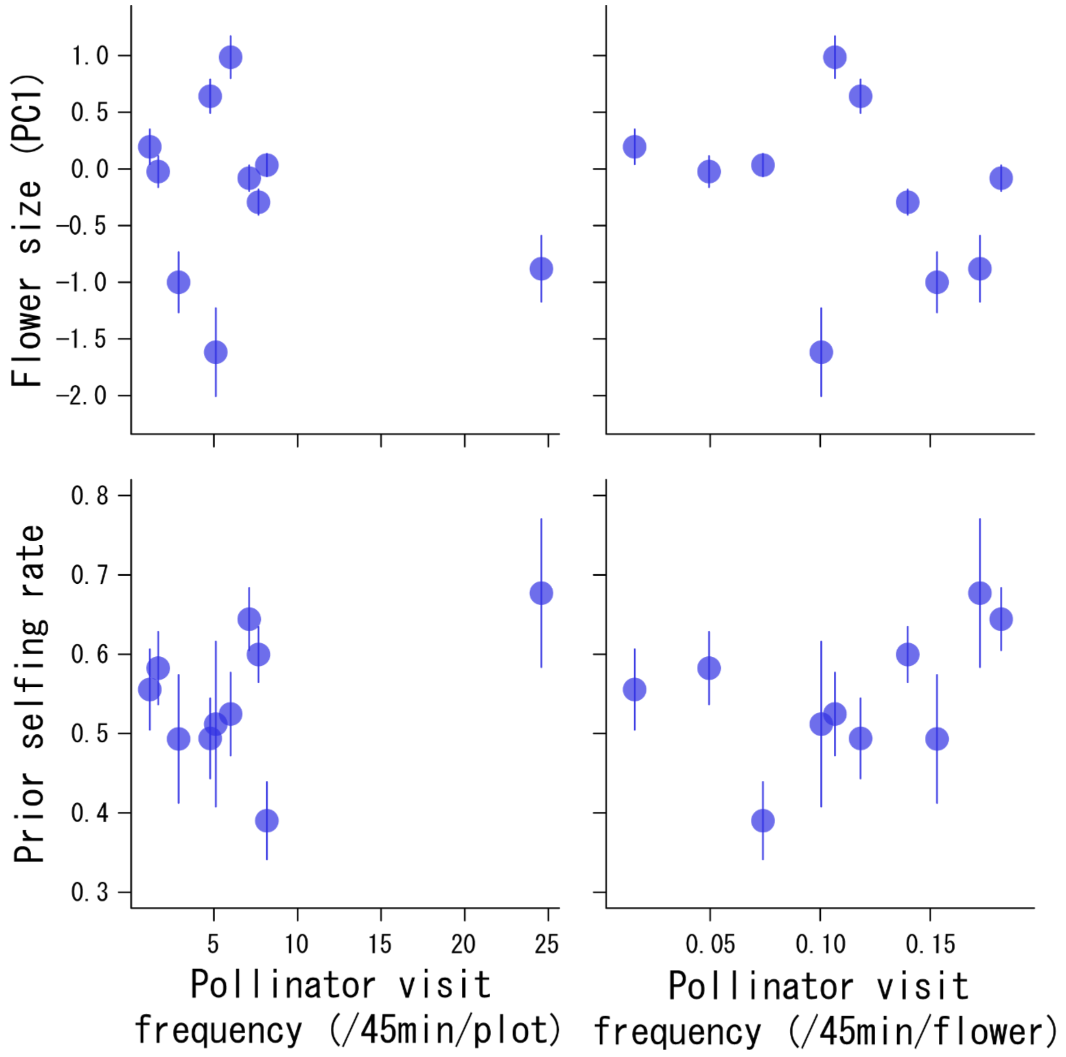

Fig S4. Relationships between pollinator visit frequency and PC1 and prior selfing rate (mean ± SE). Three 1x1 $m^2$ plots per population were examined for 45 min (15 min x 3 times) for pollinator visit observation. Pollinator availability was indexed by two measurements: the number of pollinator visits per plot for 45 minutes and the number of visits per flower for 45 minutes.

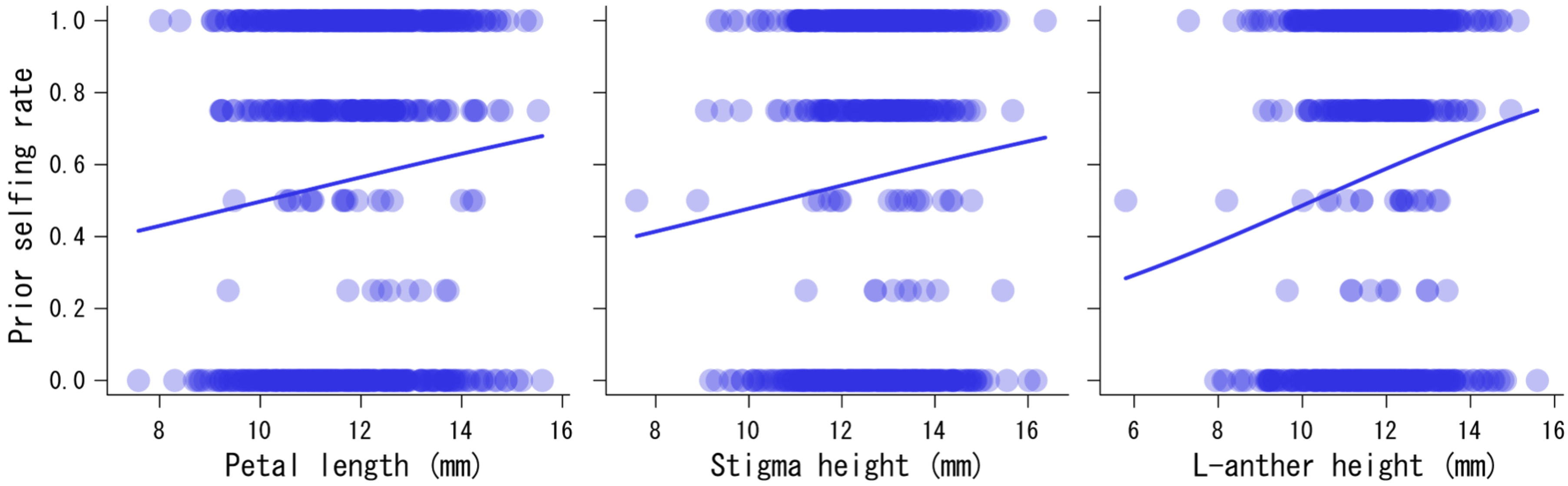

Fig S5. Relationships between the prior selfing rate and three floral traits (petal length, stigma height and L-anther height). Significant relationships are indicated with regression lines drawn using estimated coefficients from the GLMMs.

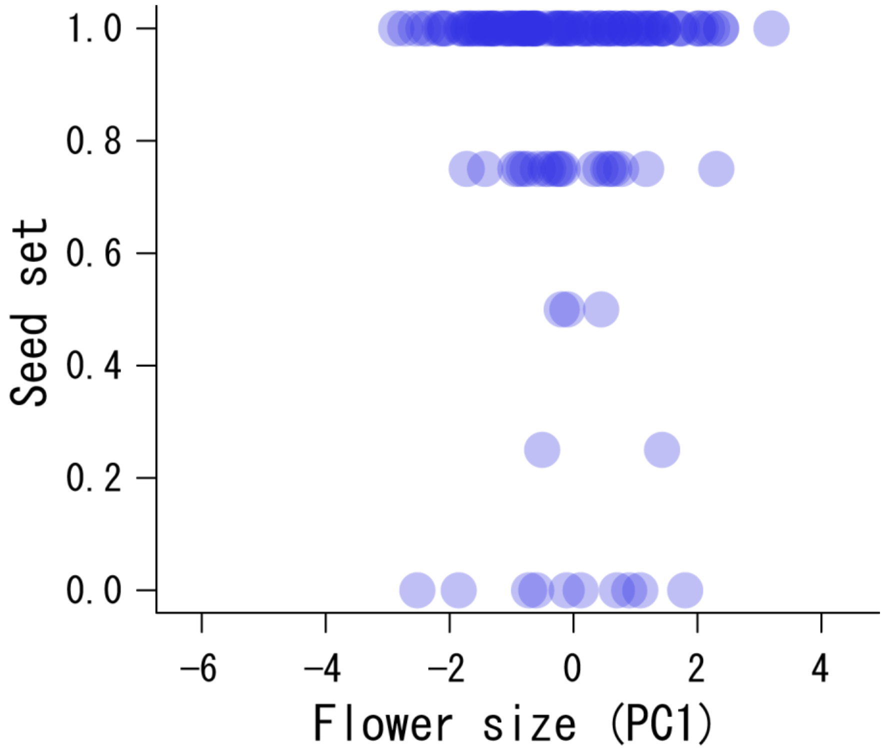

Fig S6. Relationship between flower size (PC1) and the seed set in artificially self-pollinated flowers. No significant relationship was found between the variables.

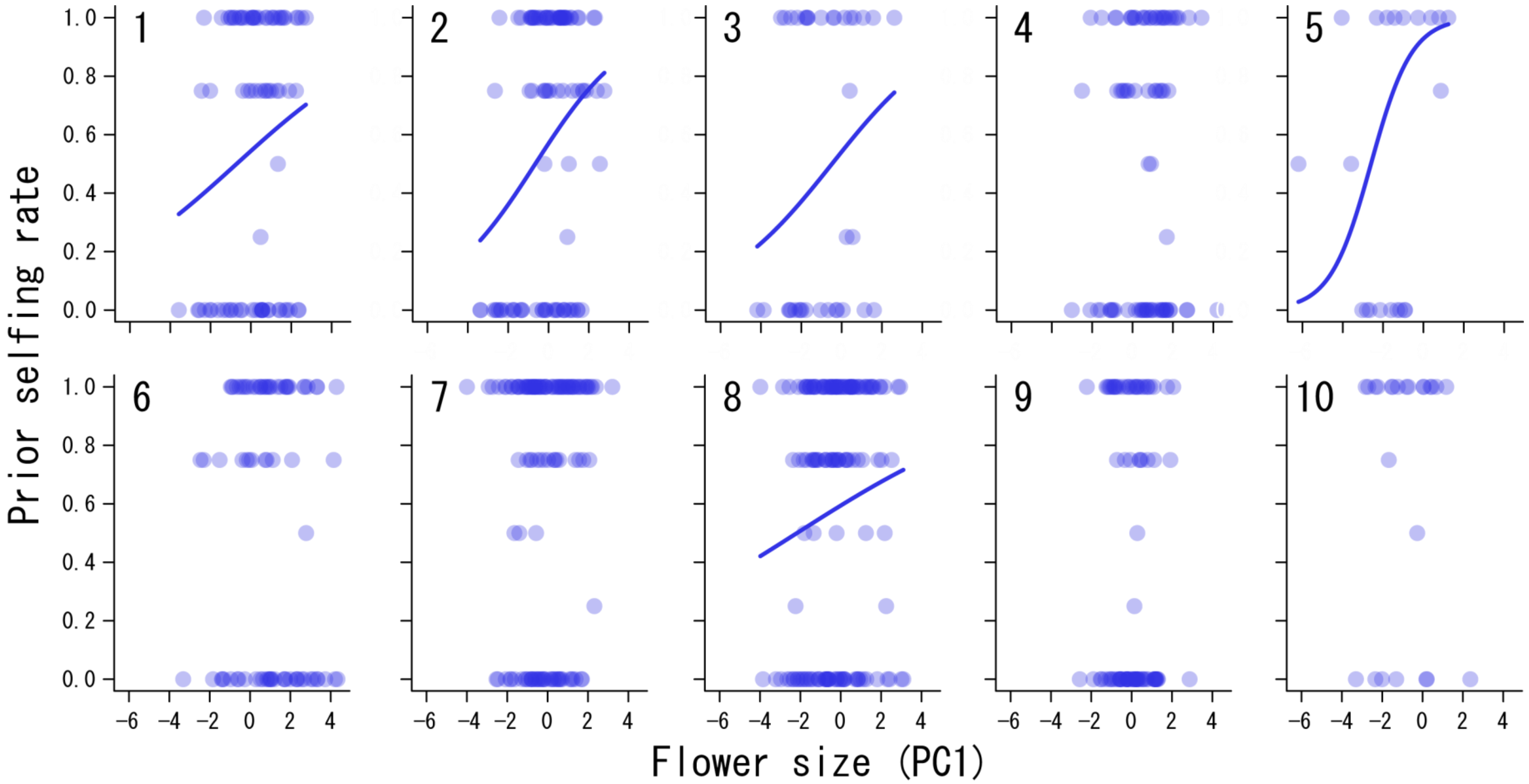

Fig S7. Relationships between flower size (PC1) and prior selfing rate for each population. Significant relationships are indicated with regression line drawns using estimated coefficients from GLMM.

Appendix S2. An experiment to timing and mechanism of prior self-pollination.

**Methods**

In 2020, to examine timing and mechanism of prior self-pollination, we observed 80 flowers from six populations before and after opening to check the occurrence of anther dehiscence just before opening and the occurrence of self-pollen deposition on the stigma surface after opening. We very carefully opened buds to avoid artificial accidental pollen deposition on the stigmas. The relationship between the anther dehiscence before opening and prior self-pollination was examined using a GLMM (with binomial error and logit link) in which the occurrence/absence of anther dehiscence ca. 10 min. before flower opening (1/0) and the occurrence/absence of autonomous self-pollination after opening (1/0) were the explanatory and response variables, respectively and the observation date identity was the random term. Data for six populations were pooled for the analysis.

**Results**

The anther dehiscence before opening occurred in 35 flowers, out of which 27 and eight flowers were autonomously self-pollinated and unpollinated, respectively. Out of 45 flowers without the dehisced anthers before opening, and were prior self-pollination, 2 and 43 flowers were autonomously self-pollinated and unpollinated after opening, respectively. Thus, prior selfing occurred significantly more in the flowers with the anther dehiscence before flower opening (Appendix S1: Table S3).

Appendix S3. An experiment to examine speed of self-pollen tube growth to the ovary.

**Methods**

In 2021, we examined a style cut experiment to examine speed of pollen tube growth to the ovary. We selected perfect flowers in which prior selfing occurred from 70 individuals of seven populations. We cut the base of the style (the part just above the ovary) at twelve different timing (0.5, 1, 1.5, 2, 2.5, 3, 3.5, 4, 4.5, 5, 5.5, 6 hours after flower opening). We checked fruit set (fruit initiation) several days after the treatment for each flower.

**Results**

The later the cut timing, the higher fruit set via prior selfing in the experimental flowers (Fig. S1).

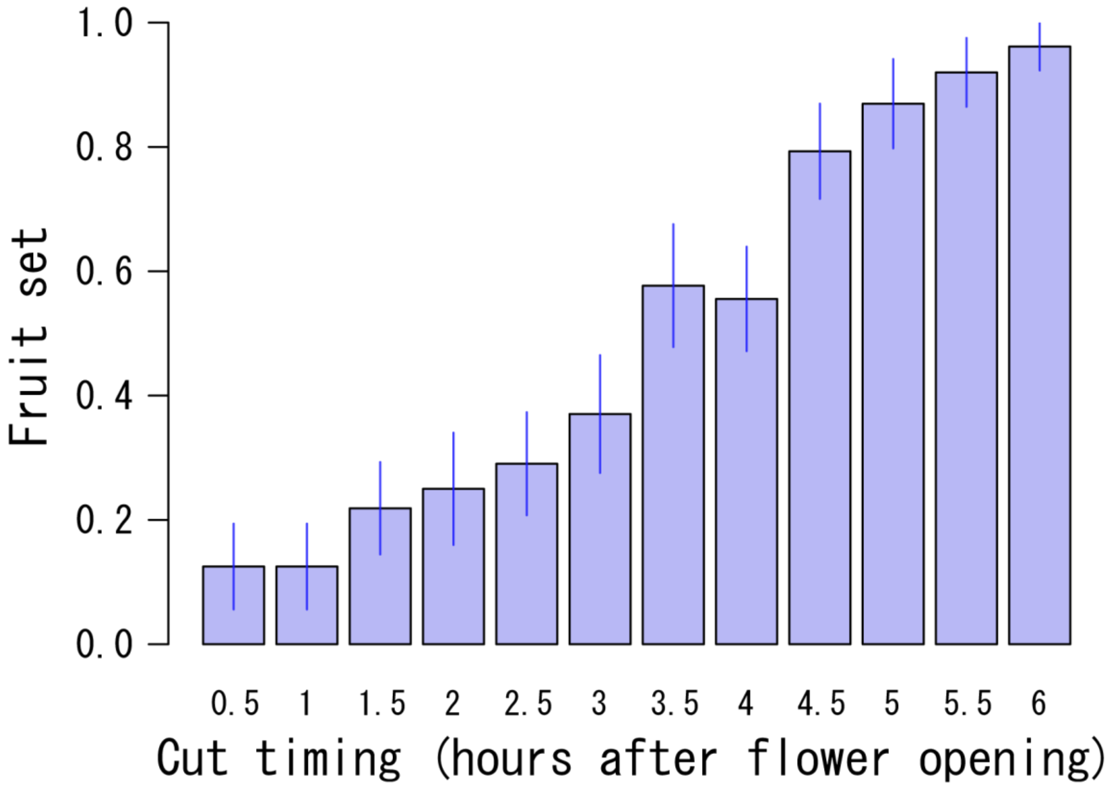

Fig S1. Fruit sets of flowers whose styles were cut at various timing after flower opening.
